# A Prefrontal Cortex-Nucleus Accumbens Circuit Attenuates Cocaine-conditioned Place Preference Memories

**DOI:** 10.1101/2025.03.21.644656

**Authors:** Xiaobo Wu, Aya M. Kobeissi, Hannah L. Phillips, Huihui Dai, Wei-Dong Yao

## Abstract

The infralimbic (IL) subregion of the prefrontal cortex (PFC), via its descending projection to the nucleus accumbens (NAc), inhibits cue-induced drug seeking and reinstatement, but the underlying mechanisms are not fully understood. Here we show that the intrinsic membrane excitability of IL layer 5 pyramidal neurons projecting to the NAc shell (IL-NAcSh neurons) suppresses cocaine-associated memories. Following repeated cocaine exposures in a conditioned place preference paradigm, IL-NAcSh neurons anatomically traced by fluorescent retrobeads undergo prolonged decrease of membrane excitability, lasting for at least 15 days after cocaine withdrawal. This persistent IL-NAcSh neuron hypoexcitability was accompanied by an increase in the rheobase, an increase in the afterhyperpolarization potential, and a decrease in the membrane input resistance. This cocaine induced neuroadapation in intrinsic excitability was not observed in prelimibic cortex neurons projecting to the NAc core (PL-NAcCo neurons), a separate descending circuit thought to promote cue-triggered drug seeking. Chemogenetic restoration of IL-NAcSh neuron activity extinguishes both the acquisition and retention of cocaine conditioned place preference memories. Our results provide direct support for the notion that the IL-NAcSh circuit serves to extinct drug associated memories and restoring the drug impaired excitability of IL-NAcSh neurons has the potential to mitigate drug-cue association memories and drug seeking.

## Introduction

Addiction is characterized by a loss of behavioral control over drug taking despite adverse consequences. Accumulating evidence from human brain imaging and animal studies suggest that dysfunction of the prefrontal cortex (PFC) leads to loss of inhibitory control and executive impairments and promotes compulsive drug use [1–3]. The infralimbic (IL) region of the medial PFC (mPFC) is of particular importance as it is involved in extinction and inhibition of drug seeking, suggesting an anti-relapse function [4–8]. In contrast, the prelimbic (PL) cortex, an adjacent more dorsal subregion of the mPFC is believed to support active reward seeking behaviors [5, 7], although an absolute PL-IL dichotomy is unlikely [9]. The IL preferentially projects to the nucleus accumbens (NAc) shell (NAcSh) [10], and the IL-NAcSh projection normally inhibits cue-induced cocaine seeking [4, 11–15] and undergoes synaptic remodeling following cocaine experience [11, 16]. Neurons in the IL, with its NAc projections, encode information about drug-associated cues and motivation for drug seeking [9, 15, 17, 18] and are hypothesized to mediate extinction of drug memories that perpetuate relapse [5]. Although the normal role of the IL-NAc circuit in inhibiting drug seeking following extinction are generally well recognized, the underlying mechanisms remain incompletely understood, and in particular, how the IL-NAc circuit regulates drug memories has not been directly investigated.

A cornerstone theory for the neurobiology of addiction posits that drug experiences elicit neuroadaptations in dopamine reward circuits, which in turn mediate addictive behaviors [19, 20]. An important player in addiction is drug-induced intrinsic plasticity in reward circuits, including the PFC [21, 22]. A number of studies have investigated intrinsic neuroadaptations of PL neurons following chronic in vivo cocaine experiences with or without withdrawal/extinction, which generally reveals an increase in excitability of PL deep layer neurons [23–27] (but see [3]). In contrast, limited studies done in the IL have found reduced neuronal excitability following cocaine experiences [15, 25]. In either mPFC subregion, circuit-specific intrinsic plasticity induced by cocaine experiences has not been examined. Of note, a recent study using cellular resolution Ca2+ imaging demonstrates that cocaine seeking behavior leads to a decrease in the activity of IL-NAc projection pyramidal neurons in rats [15]. However, neither the underlying mechanism nor the nature of cocaine-induced IL hypoexcitability was investigated.

In this study, we investigated how in vivo cocaine experience affects the intrinsic properties of pyramidal projection neurons in specific mPFC-NAc circuits. We used the cocaine conditioned place preference paradigm (CPP), a drug-environment cue associative memory model. We show that IL-NAcSh, but not PL-NAcCo, circuit displays long-lasting reduction in intrinsic excitability following withdrawal from cocaine CPP, and that normalizing IL-NAcSh circuit activity is sufficient to extinguish cocaine-conditioned place preference memory. Our findings provide insights about circuit specific neuroadaptations in the PFC following cocaine experiences and identify an important role for the IL-NAcSh circuit in regulating drug-cue association memories.

## Materials and Methods

### Animals

Male and female C57BL/6 mice (ages 4-16 weeks) were used in this study. Mice were housed 2-5 per cage with *ad litbitum* access to standard lab chow and water under a 14/10-hour light/dark cycle. The housing environment was maintained at 21-23 °C and 40-70% humidity. All studies were conducted in accordance with the guidelines of the National Institutes of Health Guide for the Care and Use of Laboratory Animals approved by the SUNY Upstate Medical University institutional animal care and use committee.

### Retrobeads and viral injections

Stereotaxic surgeries were performed as we described previously [28]. Mice were deeply anesthetized with isoflurane (5% induction, 1.5% maintenance) and head-fixed on a stereotaxic apparatus (Kopf Instruments). Bilateral burr holes were drilled with a dental drill using the following coordinates (relative to bregma): anterior/posterior [AP] +1.53 mm, medial/lateral [ML] ±0.8 mm). A 32 G 1.0 μl Neuros syringe (Hamilton Company) was carefully lowered into the NAc (depth: −4.82 mm). A volume of 80-120 nl green or red Retrobeads (Lumafluor) was injected over a period of 3 min. For retro-DREADD behavioral rescue experiment, rAAV2-EF1α-mCherry-IRES-WGA-Cre (University of North Carolina (UNC) Viral Vector Core, Chapel Hill, NC; 1 x 10^13^ vg/ml; 400 nl/side at 50 nl/min) was microinjected bilaterally into the NAc (coordinates: [AP] +1.10 to +1.54 mm, [ML] ±0.35 to ±0.58 mm, DV −3.2 to −4.0 mm), the Cre-dependent Gq-DREADD (AAV2-hSyn-DIO-hM3D(Gq)-mCherry (Addgene, Watertown, MA; 5.1-6.1 × 10^12^ vg/ml; 500 nl/side at 50 nl/min) was injected into the PFC (coordinates: [AP] +1.92 to +1.94 mm, [ML] ±0.35 to ±0.375 mm, [DV] −2.5 to −3.2 mm). Before and after injections, the syringe remained in place for 5 minutes for complete viral delivery. The incision was sutured with VetBond® Tissue Adhesive (3M). Buprenorphine (0.1 mg/kg) was injected subcutaneous as a post-operative analgesic. Mice were allowed to recover for 10-14 days before use in behavior experiments. Injection sites were confirmed postmortem following behavioral testing.

### Histology

7 to 10 days after retrobeads injections or 2-10 weeks after AAV injections, mice were sacrificed and brains were fixed with 4% v/v paraformaldehyde overnight. 150 µm-thick brain sections contained PFC and NAc regions were cut with a vibratome, rinsed in PBS, and mounted on slides. Images were taken with an Olympus fluorescent microscope.

### Behavior

Cocaine conditioned place preference (CPP) was performed as previously described with moderate changes [29, 30]. Mice were handled for 1-2 days and then received sham intraperitoneal (i.p.) injections consisting of 0.9% saline for 3-5 days. Behavior was measured in a place preference apparatus consisting of one neutral compartment enclosed by two manual guillotine doors and two (15 cm × 15 cm) conditioning chambers (MedAssociates). Each of the conditioning chambers had distinct visual (white and black walls), floor texture (grid or mesh), and olfactory cues (cinnamon or vanilla). For baseline tests, mice were placed into a neutral area and allowed free access to explore the entire apparatus for 18 min. Mice were excluded from subsequent experiments if they had a preference (> 75%) for either one of the test chambers. During the cocaine conditioning phase, a drug-paired chamber was either randomly assigned in a counter-balanced fashion or assigned to the least preferred chamber during the baseline test for each mouse. Mice received an i.p. injection of 20 mg/kg cocaine (Sigma-Aldrich) dissolved in 0.9% saline or saline on alternating days and were placed in the respective chamber for 25 minutes, for a total of 10 days. CPP tests were performed 1 (withdrawal day 1 (WD1) or 15 days (WD15) after the last cocaine conditioning session where the mice were allowed to explore the entire apparatus for 18 minutes. The place preference score (PPS) was calculated as the time in the cocaine injected side minus the time in the saline injected side. After each behavior test, mice were immediately (< 15 min) sacrificed for electrophysiology or returned to their home cages for 15-20 days (15 days withdrawal; WD15). After WD15, the mice were subjected to the CPP test again and then sacrificed for electrophysiology. For chemogenetic rescue experiments, a CPP test was conducted in a similar manner with the following modifications: on WD1, mice received CPP testing with no injection to measure acquisition of CPP behavior. On WD2 and WD15, one-half of the mice received 0.9% saline and the other half received 1 mg/kg clozapine-N-oxide (CNO) (i.p.) 20 minutes before the start of CPP tests, counter-balanced.

### Slice electrophysiology

PFC slice preparation and patch-clamp recordings were conducted as described previously [31–33]. Briefly, a mouse was sacrificed and the brain quickly removed from the skull. The brain was immediately immersed in ice-cold artificial cerebrospinal fluid (aCSF) pre-bubbled with 95% O_2_ and 5% CO_2_. Sucrose-rich aCSF contains (in mM): 235 sucrose, 25 NaHCO_3_, 2.5 KCl, 1.25 NaH_2_PO_4_, 1 CaCl_2_, 2.5 MgCl_2_, and 10 glucose, bubbled with 95% O_2_ plus 5% CO_2_. Coronal cortical brain slices (250 μm) containing the PL and IL were cut using a vibratome (Series VT1200; Leica, Germany). Slices were incubated with oxygenated aCSF containing (in mM): 126 NaCl, 2.5 KCl, 1.2 NaH_2_PO_4_, 2.5 CaCl_2_, 1.2 MgCl_2_, 25 NaHCO_3_, and 25 glucose at room temperature for at least 1 hr prior to electrophysiology recordings. For experiments presented in Figure 5, this standard aCSF was used for sectioning and pre-incubation before electrophysiology.

Pyramidal cells in the mPFC were visualized using a BX51WI infrared-differential interference contrast (IR-DIC) microscope (Olympus, Japan) with epifluorescence, optiMOS camera, and a cool-300 high-power LED fluorescence light source. Whole-cell patch-clamp recordings on neurons in the mPFC were made using a patch clamp amplifier (Multiclamp 700B; Molecular Devices, Sunnyvale, CA). Data acquisition and analysis were performed using a digitizer (DigiData 1500; Molecular Devices) and the analysis software Clampfit 10.7 (Molecular Devices), respectively. Pyramidal neurons in mPFC L5 were identified by their pyramidal morphology and fluorescence marked by green or red Retrobeads. Patch pipettes were filled with an intracellular solution containing the following (in mM): 142 KCl, 8 NaCl, 10 HEPES, 0.4 EGTA, 2 Mg-ATP, 0.25 GTP-Tris, pH 7.2-7.4 (adjusted by KOH). The GABA_A_ receptor antagonist picrotoxin (PTX, 50 μM) and AMPA receptor antagonist CNQX (20 μM) were included in the superfusion medium throughout the experiment. All recordings were made at 32°C with a temperature controller (Warner Instruments). Data were digitized at 10 kHz. The pipette resistance was 4 - 8 M when filled with intracellular solution. After a stable whole-cell patch was achieved, recordings were conducted in the current-clamp mode. Recording was done at the resting membrane potential (RMP), and a sequence of step currents ranging from −200 to +300 pA, with 25-pA increments, were applied for 500 ms each pulse, and with a 15-second interval between pulses. Passive (RMP, Input resistance, ADP, and voltage sag) and active (AP threshold, half width, rise and decay slops, peak, rheobase, and fAHP) membrane excitability properties were characterized as we described previously [28].

For validation of chemogenetic experiments, 1-month-old C57BL/6J mice injected with AAV2-hSyn-DIO-hM3D(Gq)-mCherry (Addgene) in the IL and rAAV2-EF1α-mCherry-IRES-WGA-Cre (UNC Viral Vector Core) in the NAc shell were sacrificed 2-8 weeks post injection for ex-vivo slice electrophysiology. Pyramidal neurons in the infralimbic regions expressing hM3D(Gq)-mCherry were identified based on fluorescence and morphology using IR-DIC microscopy (Olympus). Whole-cell recordings were performed as above. Resting membrane potential and action potential firing were recorded in current-clamp configuration in aCSF and then repeated with 5 μM clozapine-N-oxide (CNO freebase, HelloBio) applied to the bath.

### Statistical Analysis

Statistical tests were performed using SPSS 13.0 or GraphPad Prism (8.0) software. Differences in electrophysiological parameters in two groups were analyzed using Student’s t-test. The Mann-Whitney non-parametric test was used for data sets not passing a normal distribution. The firing number and frequency adaptation were analyzed using ANOVA followed by Fisher’s least significant difference (LDS) post hoc tests. PPS scores in figure 2 were analyzed by Student’s t-tests or Mann-Whitney non-parametric test. PPS scores in figure 5 were analyzed by ANOVA with Tukey’s post hoc test and Student’s t-test.

## Results

### Intrinsic properties of infralimbic layer 5 pyramidal neurons projecting to the NAc shell

We first investigated the intrinsic membrane properties of IL layer 5 (L5) pyramidal cells that project to the NAc shell (IL-NAcSH L5 PNs) because these projection-specific neurons have not been characterized in mice. We injected the retrograde tracer Retrobeads into the NAc shell (Fig. 1a) and identified red fluorescent Retrobeads-containing neurons in the IL (Fig 1b). Intrinsic membrane properties of IL-NAcSh L5 PNs were measured using whole-cell patch-clamp electrophysiology (Fig 1c). IL-NAcSh L5 PNs had a resting membrane potential (RMP) of −64.99 ± 1.07 mV and an input resistance of 250.89 ± 27.80 MΩ (Table 1). In response to depolarizing current injections, IL-NAcSh L5 PNs fired action potential (AP) spikes with characteristic amplitude, threshold, waveform, and adaptive frequency response (Table 1, Fig. 1d-g; Fig S1a1-b1). Prominent fast hyperpolarizing potentials (fAHP; 6.36 ± 0.72 mV; Fig. 1d; Table 1) immediately following an AP were seen, supporting the likely presence of voltage-gated Kv7 K+ channels and Ca2+ activated K+ channels that contribute to AHPs [27, 34, 35]. IL-NAcSh L5 PNs also displayed prominent depolarizing sag (6.27 ± 0.76 mV; Table 1) and afterdepolarization potentials (ADPs, 6.34 ± 0.66 mV; Table 1) in response to hyperpolarizing currents (Fig. 1h), indicating activation of h-currents (I_h_, hyperpolarization-activated nonspecific cation current). I_h_ is mediated by hyperpolarization-activated cyclic nucleotide gated channel (HCN) channels, a canonical feature of most L5 PNs [36] that contributes to voltage sags and ADPs and modulates intrinsic excitability [37, 38]. Together, these results define the passive and active membrane properties of mouse IL-NAcSh L5 PNs and are consistent with intrinsic membrane properties of rodent IL neurons in previous studies [39, 40].

**Figure 1.**
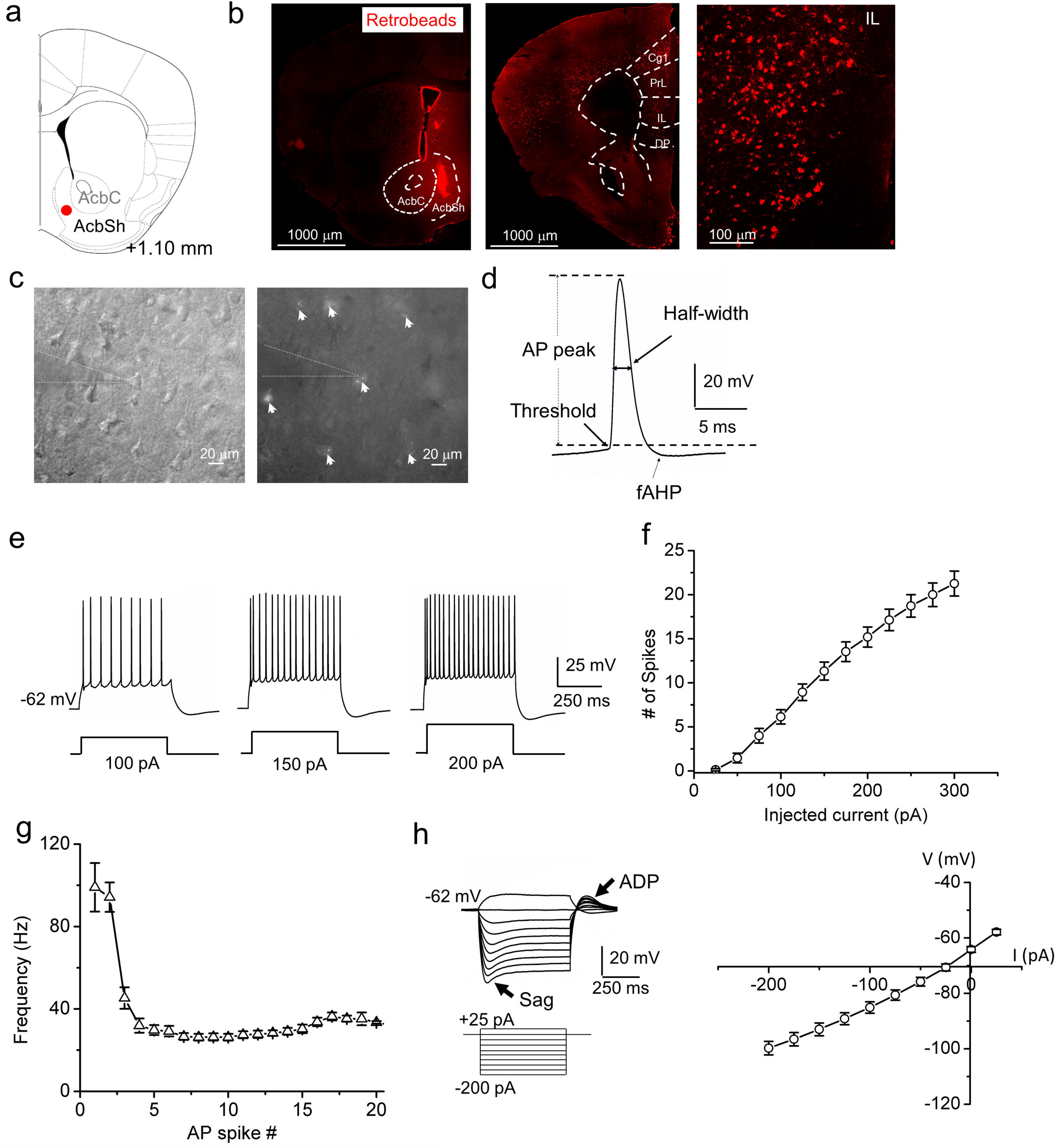
Electrophysiological characterization of IL-NAcSh L5 pyramidal neurons. (**a**) Schematic of retrobead injection site in the NAc shell. **(b)** Fluorescence images of retrobeads injected at the NAc shell (left) and retrogradely traveled to mPFC and other cortical and subcortical regions at low (middle) or high (right) magnifications. (**c**) DIC (left) and fluorescent (right) images of an mPFC slice showing patch-clamp recording of a retrobeads-labeled layer 5 pyramidal neuron in the IL. White arrows indicate retrobeads-positive neurons. (d) Schematic of action potential (AP) characterizations. scale bars: 20 mV, 5 ms. (**e**) Representative recordings of AP spikes in response to 100, 150 and 200 pA step current injections in IL-NAcSh L5 PNs. Scale bars: 25 mV and 250 ms. (**f**) AP spike number vs. current injection relationship in IL-NAcSh L5 PNs. N = 15 cells from 6 mice. (**g**) Summary spike frequency adaptation in response to 200 pA step current injection. N = 15 cells from 6 mice. (**h**) Representative recording of membrane voltages in response to sub-threshold depolarizing or hyperpolarizing currents (left) and summary current-voltage relationship (right) in IL-NAcSh L5 PNs. Scale bars: 20 mV, 250 ms. N = 15 cells from 6 mice.

**Table 1.** Passive and active membrane properties of IL-NAcSh and PL-NAcCo L5 pyramidal neurons.

|  | IL-NacSh neurons<br>(N = 15) | PL-NacCo neurons<br>(N = 16) |
| --- | --- | --- |
| RMP (mV) | -64.99 ± 1.07<sup>**</sup> | -69.71 ± 0.87 |
| V<sub>th</sub> (mV) | -42.17 ± 1.10 | -42.82 ± 1.03 |
| Input Resistance (MΩ) | 250.89 ± 27.80 | 310.49 ± 26.5 |
| Half-width AP (ms) | 1.25 ± 0.37 | 1.29 ± 0.05 |
| fAHP (mV) | 6.36 ± 0.72 | 5.99 ± 0.82 |
| ADP (mV) | 6.34 ± 0.66<sup>***</sup> | 2.45 ± 0.3 |
| Sag (mV) | 6.27 ± 0.76<sup>***</sup> | 2.71 ± 0.33 |
| AP peak (mV) | 79.52 ± 2.98 | 77.67 ± 3.67 |
| Rheobase (pA) | 66.67 ± 5.27 | 75.00 ± 7.57 |
| Rise slope (mV/ms) | 162.04 ± 12.71 | 157.38 ± 14.91 |
| Decay slope (mV/ms) | 55.13 ± 3.72 | 51.39 ± 4.09 |

RMP, resting membrane potential; V<sub>th</sub>, voltage threshold; AP, action potential; fAHP, fast afterhyperpolarization; ADP, after depolarization; AP, action potential Statistical tests: Student's t-test, <sup>\*\*</sup> $P < 0.01$ , <sup>\*\*\*</sup> $P < 0.001$ .

### Prolonged hypoexcitability elicited by cocaine CPP in IL-NAcSh L5 PNs

To examine the effects of cocaine place conditioning on the intrinsic excitability of IL-NAcSH L5 PNs, we first established a cocaine CPP assay on Retrobeads-injected mice. Two weeks following stereotaxic retrobeads injection surgery, mice were subjected to a cocaine CPP regimen (Fig. 2a; modified from [29, 30]). Cocaine administration was paired to a neutral context (cocaine-paired context) and saline administration to another context with different cues (saline-paired context) (Fig. 2 b). Saline-only administered mice were used as controls. Cocaine treated mice spent significantly more time in the cocaine-paired context compared to saline treated mice both at short-term, 1-day, withdrawal (Coc WD1, 268.04 ± 19.05, Sal, 15.25 ± 38.44; *P* < 0.0001, Student’s *t*-test) and long-term, 15-day, withdrawal (Coc WD15, 224.7 ± 37.40, Sal, 26.93 ± 32.83; *P* = 0.0007, Fig. 2 c, Mann-Whitney test), indicating successful acquisition and retention of cocaine-conditioning memories.

**Figure 2.**
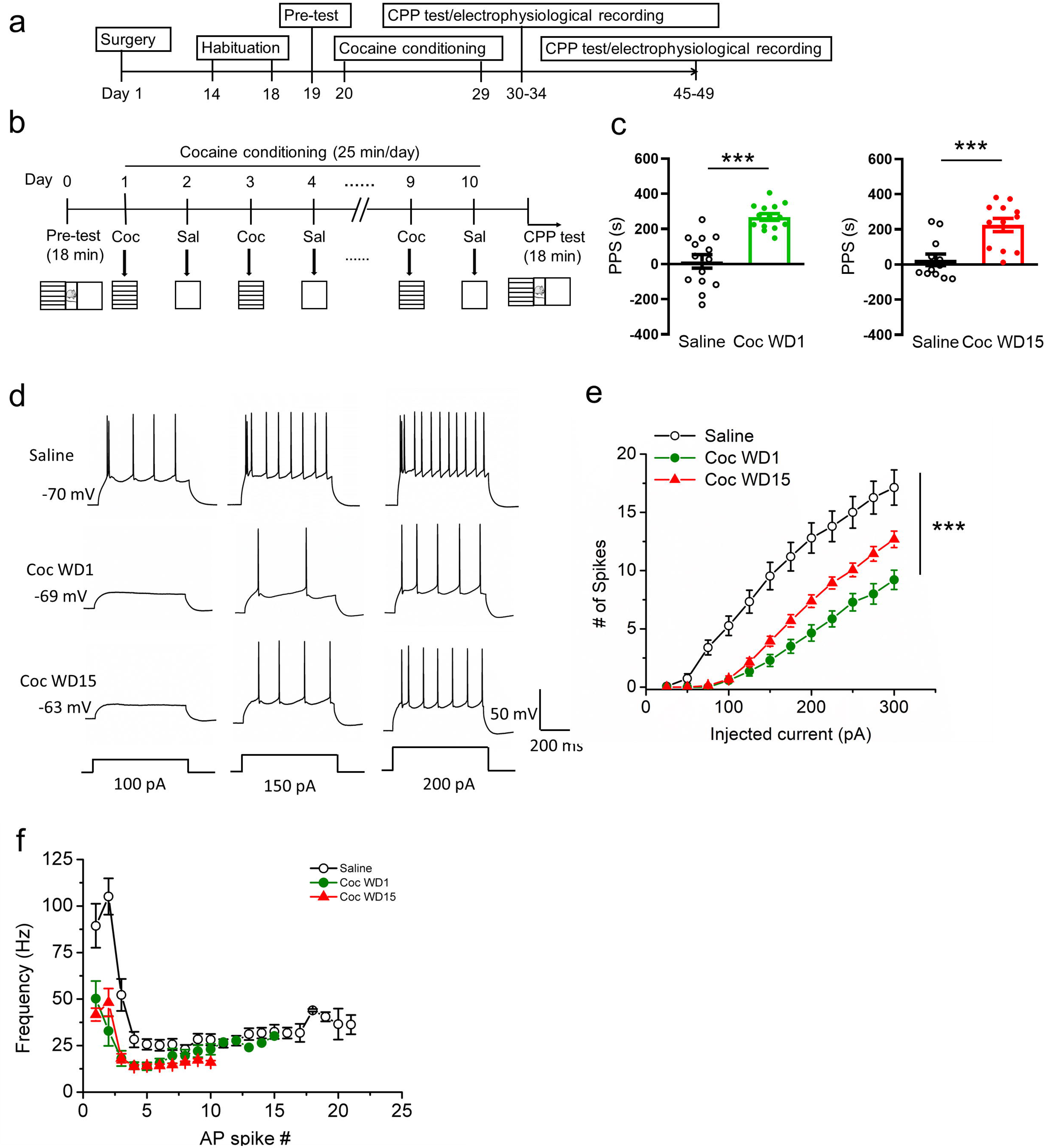
Sustained hypoexcitability and spike frequency adaptation in IL-NAcSh L5 pyramidal neurons following cocaine CPP. (**a**) Experimental timeline. (**b**) Schematic of cocaine CPP assay. (**c**) Cocaine place preference score (PPS) on withdrawal day 1 (WD1) and day 15 (WD15) following CPP. Saline vs. Coc WD1, \*\*\*\**P* < 0.0001, Student’s t-test; Saline vs. Coc WD15, \*\*\**P* < 0.001, Mann-Whitney test). N = 14 (saline vs Coc WD1) and 12 (saline vs Coc WD15) mice. (**d**) Representative AP spikes elicited by step current injections (100, 150 and 200 pA) in IL-NAcSh L5 PNs in saline, cocaine WD1 and cocaine WD15 mice. (e) Summary of AP spike numbers vs. step currents in IL-NAcSh L5 PNs. \*\**P* < 0.01, \*\*\**P* < 0.001, two-way ANOVA with RM. Saline, N = 15 cells from 7 mice; Coc WD1, N = 14 cells from 8 mice; Coc WD15, N = 16 cells from 8 mice. (**f**) Summary data of spike frequency adaptations of IL-NAcSh L5 PNs in different groups in response to a 200 pA step current injection. Saline, N = 15 cells from 7 mice; Coc WD1, N = 14 cells from 8 mice; Coc WD15, N = 16 cells from 8 mice.

Previous studies reported that in vivo cocaine exposures reduced IL neuronal activity in rats [15, 25], but neither the circuits involved nor the underlying mechanisms were investigated. To assess how cocaine CPP affects the excitability mechanisms of IL-NAcSh neurons, we characterized the passive and active membrane properties of IL-NAcSh L5 PNs following 1-day or 15-day withdrawal from CPP. On both WD1 and WD15, cocaine-exposed mice showed significantly reduced action potential spiking in response to depolarizing current injections (Fig. 2d,e), accompanied by significantly higher rheobase (the minimal currents needed to elicit an AP spike (Coc WD1, 141.07 ± 11.9 pA; Coc WD15, 120.31 ± 6.13 pA; Sal, 70.0 ± 6.07 pA; F_2,42_ = 19.36, *P* < 0.0001, one-way ANOVA with post-hoc LSD test; Table 2), fAHP (Coc WD1, 11.62 ± 0.70 mV; Coc WD15, 9.39 ± 0.57 mV; Sal, 6.32 ± 1.04 mV; F_2,42_ = 10.9, *P* = 0.0002, one-way ANOVA with post-hoc LSD test; Table 2), and decreased AP half-width (Coc WD1, 1.25 ± 0.06 ms; Coc WD15, 1.17 ± 0.04 ms; Sal, 1.44 ± 0.05 ms; F2,42 = 7.74, *P* = 0.0014, one-way ANOVA with post-hoc LSD test; Table 2). In addition, IL-NAcSh L5 PNs firing was significantly reduced in response to increasing depolarizing currents in mice with cocaine exposure at both WD1 and WD15 (F_2,42_ = 20.85, *P* < 0.0001, two-way ANOVA with repeated measures (RM) test, Fig. 2e). The spike frequency adaptation was also compromised in cocaine treated mice at both WD1 and WD15 (Fig. 2f; Fig. S2). Together, these results indicate that cocaine place preference conditioning induced persistent hypoexcitability in IL-NAcSh L5 pyramidal neurons.

**Table 2.** Comparisons of electrophysiological properties of IL-NAcSh and PL-NAcCo L5 pyramidal neurons in saline and cocaine CPP mice.

|  | IL-NAcSh neurons |  |  | PL-NAcCo neurons |  |
| --- | --- | --- | --- | --- | --- |
|  | Saline<br>(N = 15) | Coc WD1<br>(N = 14) | Coc WD15<br>(N = 16) | Saline<br>(N = 14) | Coc WD15<br>(N = 10) |
| RMP (mV) | -66.63 ± 0.90 | -68.32 ± 0.99 | -65.13 ± 0.86 | -66.93 ± 1.14 | -65.26 ± 1.23 |
| Vth (mV) | -36.00 ± 0.82 | -34.43 ± 0.78 | -34.70 ± 0.77 | -36.61 ± 1.71 | -35.58 ± 0.92 |
| Input Resistance (MΩ) | 350.6 ± 32.65 | 225.52 ± 31.03 | 164.79 ± 14.74 | 332.16 ± 18.67 | 316.17 ± 49.4 |
| Half-width AP (ms) | 1.44 ± 0.05 | 1.25 ± 0.06<sup>#</sup> | 1.17 ± 0.04<sup>###</sup> | 1.44 ± 0.07 | 1.42 ± 0.08 |
| fAHP (mV) | 6.32 ± 1.04 | 11.62 ± 0.70<sup>###</sup> | 9.39 ± 0.57<sup>#</sup> | 9.5 ± 1.53 | 11.9 ± 1.36 |
| ADP (mV) | 3.99 ± 1.07 | 2.32 ± 0.32 | 3.42 ± 0.42 | 2.18 ± 0.61 | 4.1 ± 1.0 |
| Sag (mV) | 3.21 ± 0.78 | 2.56 ± 0.35 | 3.44 ± 0.50 | 1.95 ± 0.4 | 3.91 ± 0.9<sup>*</sup> |
| AP peak (mV) | 68.03 ± 2.19 | 80.37 ± 1.77 | 80.18 ± 2.65 | 71.61 ± 3.95 | 71.86 ± 2.79 |
| Rheobase (pA) | 70.0 ± 6.07 | 141.07 ± 11.9<sup>###</sup> | 120.31 ± 6.13<sup>##</sup> | 69.64 ± 5.36 | 62.5 ± 4.17 |
| Rise slope (mV/ms) | 129.89 ± 8.00 | 167.15 ± 8.00 | 177.55 ± 10.50<sup>##</sup> | 133.46 ± 13.81 | 132.54 ± 8.19 |
| Decay slope (mV/ms) | 35.35 ± 2.43 | 58.18 ± 5.19<sup>#</sup> | 56.97 ± 3.24<sup>##</sup> | 42.19 ± 4.12 | 43.6 ± 2.92 |

RMP, resting membrane potential; Vth, voltage threshold; AP, action potential; fAHP, fast afterhyperpolarization; ADP, after depolarization. Statistical tests: One-way-ANOVAs with LDS post-hoc tests, <sup>#</sup>*P* < 0.05, <sup>##</sup>*P* < 0.01, <sup>###</sup>*P* < 0.001 for IL-NAc neurons (Saline vs. Coc WD1, Saline vs Coc WD15); Student's t-test, <sup>\*</sup>*P* < 0.05 for PL-NAc neurons (Saline vs. Coc WD15).

### Lack of persistent hypoexcitability in PL-NAcCo L5 PNs following cocaine CPP

To examine whether the cocaine-induced IL-NAcSh neuron hypoexcitability was projection-specific, we recorded PL L5 PNs projecting to the NAc core (PL-NAcCo L5 PNs). This circuit mediates drug seeking and plays essential roles in active reward behavior [5, 7], but its intrinsic plasticity following in vivo cocaine experiences has not been examined. We first labeled and characterized this mPFC circuit by injecting Retrobeads into NAc core and recorded fluorescent L5 PNs in the PL (Fig. 3a, b). Whole-cell current-clamp analyses show that PL-NAcCo L5 PNs displayed more hyperpolarized RMP (IL-NAcSh, −64.99 ± 1.07 mV; PL-NAcCo, −69.71 ± 0.87 mV, *P* = 0.002, Student’s t-test), smaller voltage sag (IL-NAcSh, 6.27 ± 0.76 mV; PL-NAcCo, 2.71 ± 0.33 mV, *P* <0.0001, Student’s t-test) and ADP (IL-NAcSh, 6.34 ± 0.66 mV; PL-NAcCo, 2.45 ± 0.3 mV, p < 0.0001, Student’s t-test), but statistically similar input resistance (IL-NAcSh, 250.89 ± 27.80 M; PL-NAcCo, 310.49 ± 26.50 M, *P* = 0.13, Student’s t-test) and AP spike waveforms compared with IL-NAcSh L5 PNs (Table 1; Fig. 3c-f). In addition, PL-NAcCo L5 PNs fire slightly fewer, but statistically indifferent numbers of AP spikes in response to increasing depolarizing currents (F_1,29_ = 3.22, *P* = 0.08, two-way ANOVA with RM test; at 200 pA of step current: IL-NAcSh, 15.2 ± 1.13; PL-NAcCo, 12.63 ± 0.97, *P* = 0.09, Student’s t-test; Fig. S1a1-a2). However, the two projections show a distinct initial spike frequency response characterized by a longer delay of first spikes in PL-NAcCo neurons (IL-NAcSh, 0.27 ± 0.02; PL-NAcCo, 0.39 ± 0.02, *P* = 0.0001, Student’s t-test; 200 pA injection; Fig. S1a2-b2). These results document circuit-specific intrinsic membrane properties for mPFC projection neurons.

**Figure 3.**
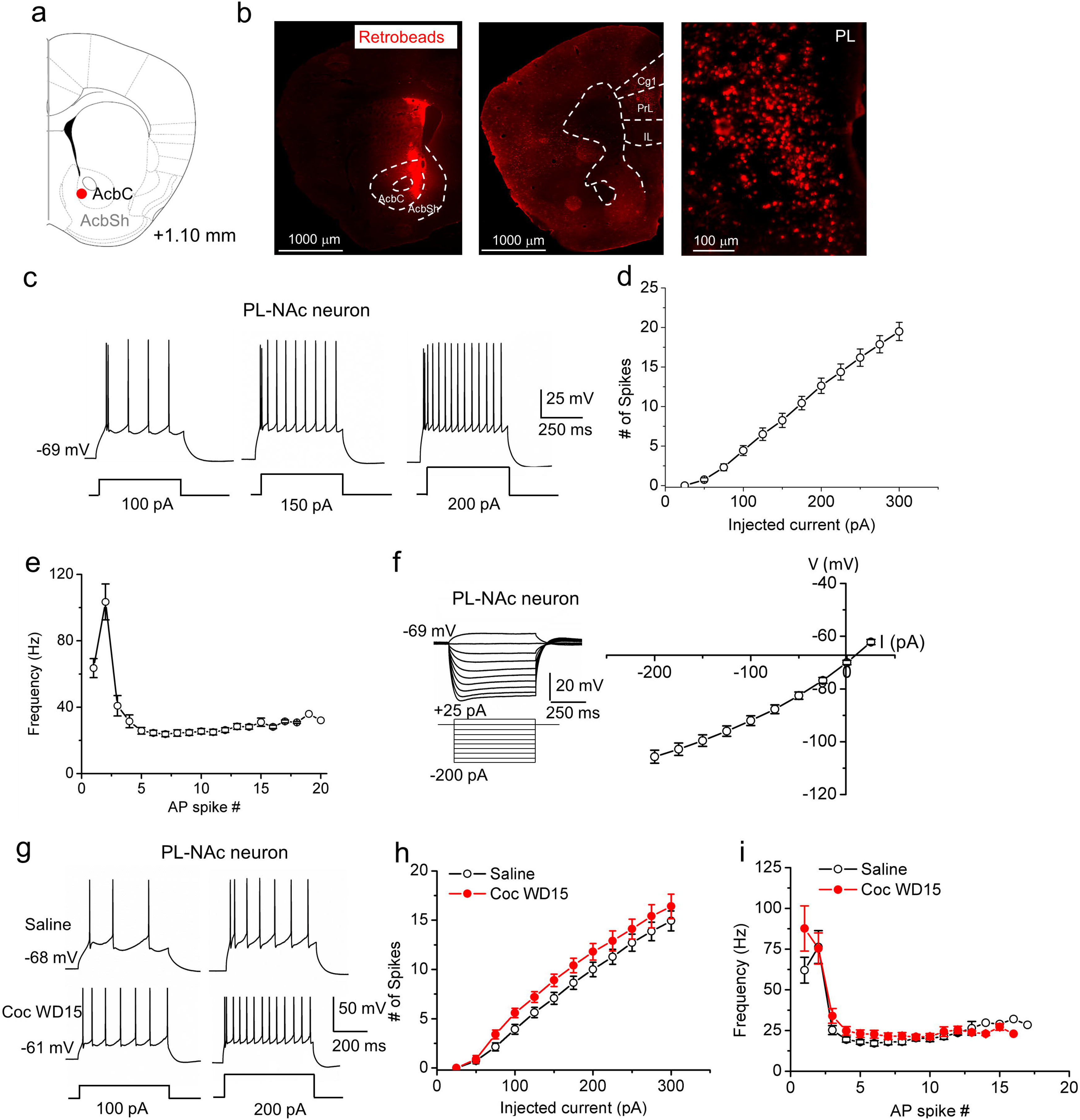
Unaltered neuronal excitability in PL-NAcCo L5 pyramidal neurons following withdrawal from cocaine CPP. (**a**) Schematic of Retrobead injection site in the NAc core. **(b)** Fluorescent images of Retrobeads injected at the NAc core (left) and traveled to the mPFC and other regions at low (middle) or high (right) magnifications. (**c**) Representative current-clamp recordings of membrane voltages in response to 100, 150 and 200 pA step current injections in PL-NAcCo L5 PNs in naive mice. (**d**) Summary AP spike number vs. current relationships in PL-NAcCo L5 PNs. N = 16 cells from 8 mice. (**e**) Summary spike frequency adaptations in response to 200 pA step current injection. N = 16 cells from 8 mice. (**f**) Representative membrane potentials in response to sub-threshold depolarizing and hyperpolarizing currents (left) and summary current-voltage relationship (right) in PL-NAcCo L5 PNs. N = 16 cells from 8 mice. (**g**) Representative AP spikes elicited by 100 and 200 pA current injections in saline and cocaine CPP mice 15 days after withdrawal. (**h**) Summary AP spike number vs. current injection relationship in PL-NAcCo L5 PNs. Saline, N = 14 cells from 6 mice; Coc WD15, N = 10 cells from 5 mice. (**i**) Summary instantaneous firing frequency in PL-NAcCo L5 PNs in response to a 200-pA step current injection in saline and WD15 cocaine CPP mice. Saline, N = 14 cells from 6 mice; Coc WD15, N = 10 cells from 5 mice..

After characterizing PL-NAcCo L5 PN intrinsic membrane properties, we examined how cocaine CPP modified PL-NAcCo L5 PN excitability. Following cocaine CPP at WD15 and compared to saline-treated mice, PL-NAcCo neurons exhibited slightly higher, but statistically insignificantly AP spikes in response to various depolarizing current injections (Fig. 3g,h; F_1,22_ = 2.26, *P* = 0.15, two-way ANOVA with RM test), statistically indistinguishable spike frequency adaptations (Coc WD15, 0.25 ± 0.03; Sal, 0.28 ± 0.02, *P* = 0.28, Student’s t-test; 200 pA injection; Fig. 3i, Fig. S3), and similar rheobase, RMP, input resistance, fAHP and AP waveforms (Table 2). These results indicate that cocaine CPP does not elicit overt lasting excitability adaptations in the PL-NAcCo circuit, supporting that cocaine experience may alter intrinsic plasticity in a circuit-dependent manner.

### Altered passive membrane properties of IL-NAcSh neurons following cocaïne CPP

To explore potential mechanisms underlying the cocaine-induced IL-NAcSh L5 PN hypoexcitability, we next characterized the passive membrane properties of these neurons by applying a series of hyperpolarizing step currents (Fig 4a). We obtained I-V curves, which showed upward shifts in WD1, which further worsened at WD 15, cocaine groups compared to saline group (Fig 4b; F_2,42_ = 9.02, *P* = 0.0006, two-way ANOVA with post-hoc LSD tests.). We measured the input resistance, which reflects the extent to which membrane channels are open at resting, by computing the slopes of I-V curves (Fig. 4c). Compared with the saline group, the input resistance was significantly decreased in IL-NAcSh L5 PNs at both WD1 and WD5 (Fig. 4c; Coc WD1, 225.52 ± 31.03 MΩ; Coc WD15, 164.79 ± 14.74 MΩ; Sal, 350.6 ± 32.65 MΩ; F_2,42_ = 12.82, *P* < 0.0001, one-way ANOVA with post-hoc LSD tests). In contrast, both the voltage sag (Coc WD1, 2.56 ± 0.35 mV; Coc WD15, 3.44 ± 0.50 mV; Sal, 3.21 ± 0.78 mV; F_2,42_ = 1.21, *P* = 0.31 one-way ANOVA with post-hoc LSD tests) and ADP (Coc WD1, 2.32 ± 0.32 mV; Coc WD15, 3.42 ± 0.42 mV; Sal, 3.99 ± 1.07 mV; F_2,42_ = 1.46, *P* = 0.24, one-way ANOVA with post-hoc LSD tests) in IL-NAcSh L5 PNs were not significantly different between saline and cocaine CPP groups (Table 2). In comparison, the input resistance of PL-NAcCo L5 PNs was not affected following cocaine CPP (Coc WD15, 316.17 ± 48.4; Sal, 332.16 ± 18.67; *P* = 0.74, Student’s t-test). Interestingly, however, the voltage sag (Coc WD15, 3.91 ± 0.9 mV; Sal,1.92 ± 0.4 mV; *P* = 0.04, Student’s t-test), and to a lesser degree ADP (Coc WD15, 4.1 ± 1.0 mV; Sal, 2.18 ± 0.61 mV; *P* = 0.09, Student’s t-test), of PL-NAcCo L5 PNs was somewhat increased in cocaine CPP mice compared to saline control (Table 2).

**Figure 4.**
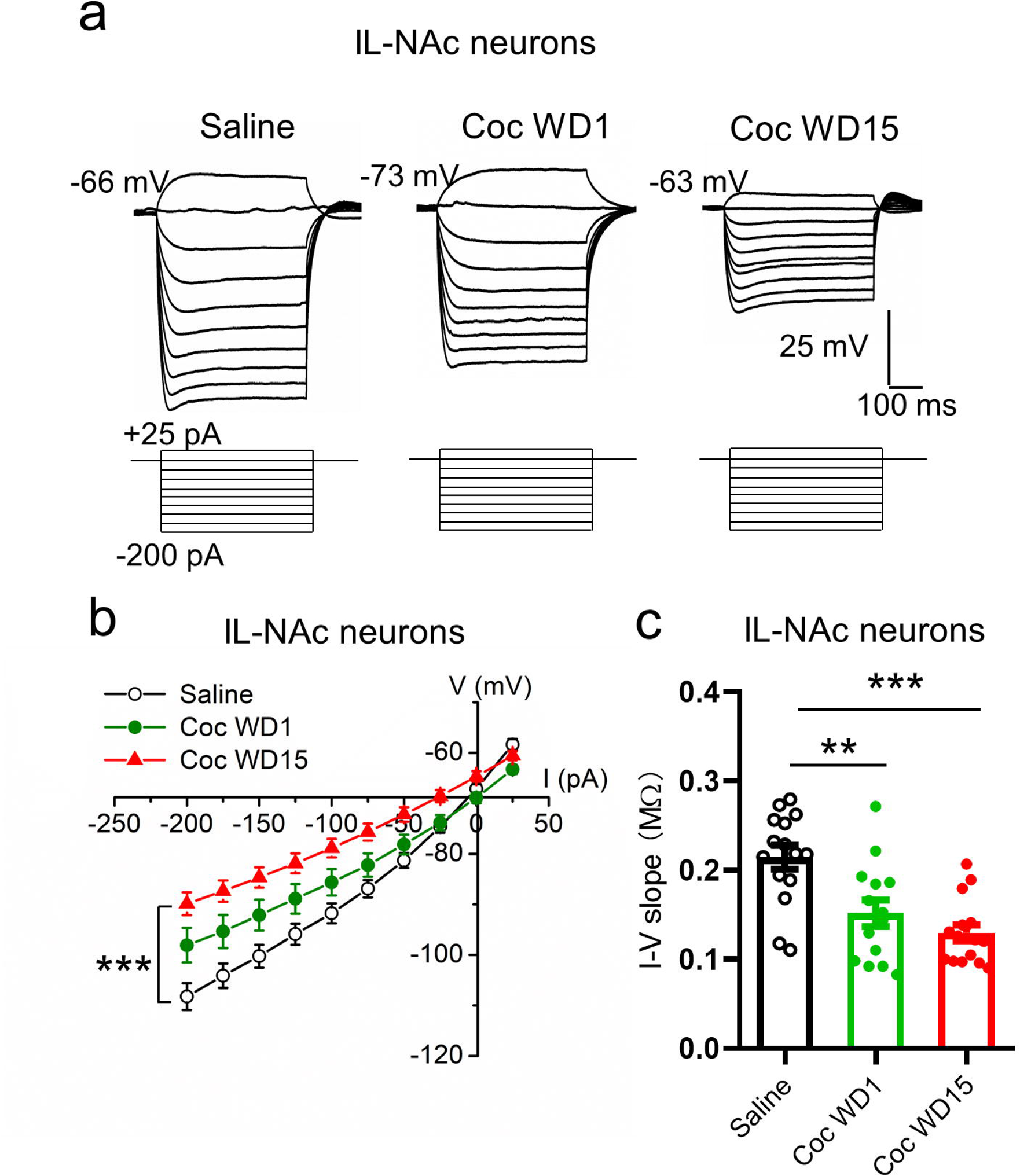
Reduced membrane input resistance in IL-NAcSh L5 pyramidal neurons following cocaine CPP. (**a**) Representative membrane potentials in response to subthreshold depolarizing and hyperpolarizing current injections in IL-NAcSh L5 PNs in saline, cocaine WD1 and cocaine WD15 mice. (**b**) Summary I-V curves of IL-NAcSh L5 PNs for various groups. *** *P* < 0.001, one-way ANOVA followed by LSD tests. (**c**) Summary I-V slopes for various groups. \*\**P* < 0.01, \*\*\**P* < 0.001, one-way ANOVA followed by LSD tests. Saline, N = 15 cells from 7 mice; Coc WD1, N = 14 cells from 8 mice; Coc WD15, N = 16 cells from 8 mice.

### Chemogenetic activation of IL-NAcSh projection attenuates cocaine conditioned memory

Finally, we examined whether the reduced IL-NAcSh circuit activity was responsible for the acquisition and retention of the cocaine-conditioned place preference memory using a chemogenetics approach (Fig. 5). To selectively activate IL-NAcSh projection neurons, mice were injected bilaterally with AAV2-hSyn-DIO-hM3d(Gq)-mCherry in the IL and rAAV2-EF1α-mCherry-IRES-WGA-Cre in the NAc shell (Fig. 5 a-c), recovered for 2 to 3 weeks, and subjected to cocaine CPP. IL-NAcSh neurons were labeled by the expression of Cre-dependent hM3d(Gq)-mCherry in the presence of Cre fused to the transsynaptic tracer WGA (WGA-Cre) from the NAc. Infected IL-NAcSh neurons expressing hM3d(Gq)-mCherry, a stimulatory designer receptor exclusively activated by designer drug (DREADD), was activated by the synthetic ligand clozapine-N-oxide (CNO). Whole-cell patch-clamp recordings on slices confirmed that bath applied CNO enhanced the excitability of hM3d(Gq)-mCherry-expressing IL-NAcSh neurons by depolarizing the RMP (vehicle, −75.06 ± 1.87; CNO, −62.29 ± 2.68; *P* = 0.007, Paired Student’s t-test), decreasing the rheobase (vehicle, 89.29 ± 14.29; CNO, 35.71 ± 7.43; *P* = 0.008, Paired Student’s t-test), and increasing AP spike numbers in response to depolarizing currents (Fig. 5d-g). Consistent with an increased excitability, some neurons were quiescent at resting but showed spontaneous firing following CNO application (Fig. 5d).

**Figure 5.**
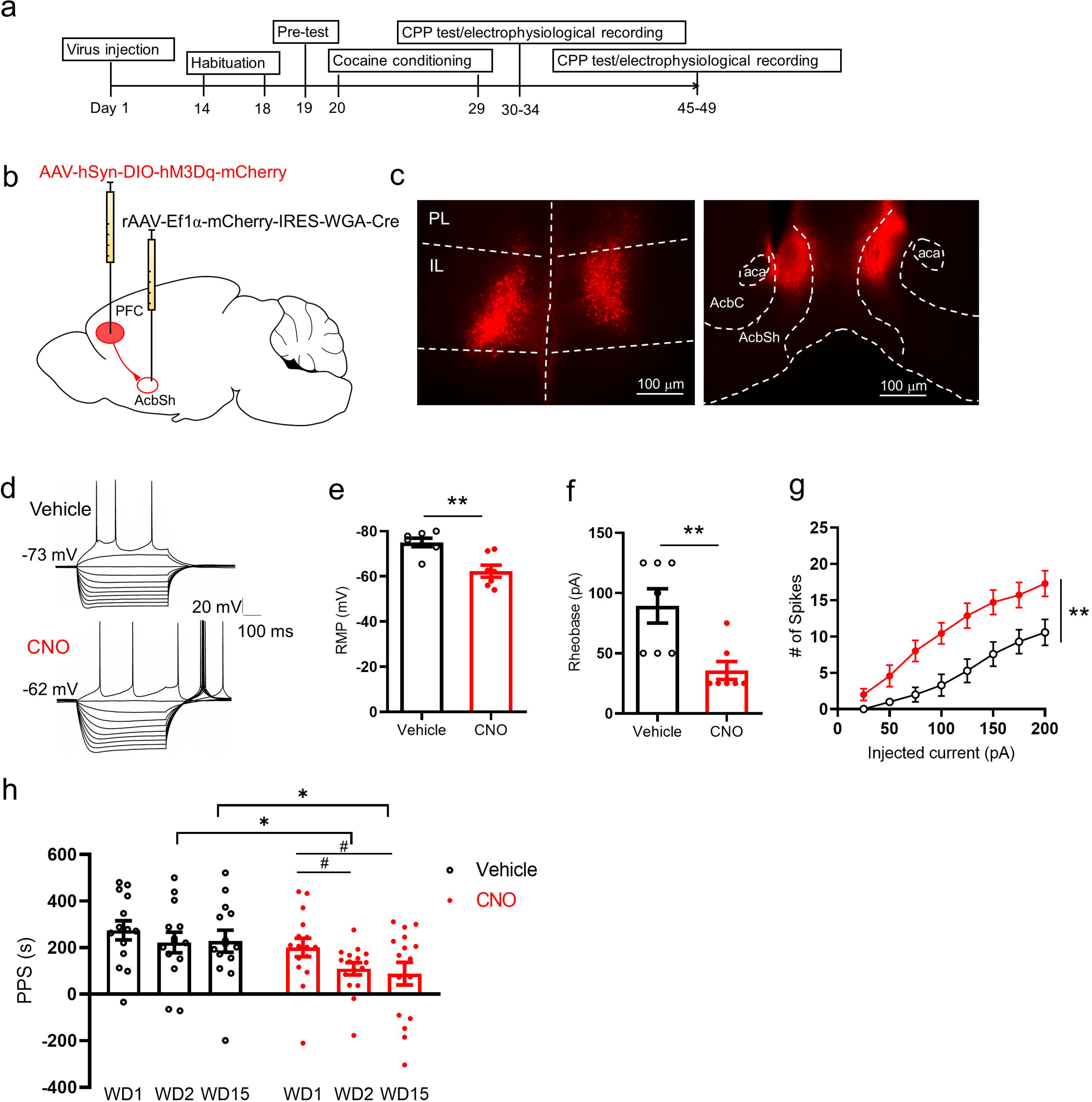
Chemogenetic activation of IL-NAcSh projection neurons attenuates cocaine place memories. (**a**) Experimental timeline. (**b**) Stereotaxic virus injection schematic. (**c**) Representative images of AAV2-hSyn-DIO-hM3D(Gq)-mCherry expression in the IL (left) and rAAV2-EF1a-mCherry-IRES-WGA-Cre expression in NAc shell (right). (**d**) Example current-clamp records showing current injection evoked AP firing in a viral transduced neuron in the IL before and after bath application of CNO (5 μM). (**e-g**) Summary of resting membrane potential (**e**), rheobase (**f**), and AP spike number-current injection relationship (**g**) before and after CNO perfusion. Statistics: **e** and **f**, \*\**P* < 0.01, Student’s *t*-tests; **g**, F_1,12_ = 9.84, \**P* < 0.01, two-way ANOVA with RM, N = 7 cells from 3 mice. (**h**) Systemic injection of CNO (1.0 mg/kg, i.p.) significantly decreased cocaine CPP at WD2 and WD15 compared to vehicle-injected controls. \**P* < 0.05, two-tailed unpaired Student’s t-test; # *P* < 0.05, one-way repeated measures ANOVA followed by Tukey’s post hoc test. N= 14 mice (vehicle group) and N= 16 mice (CNO group).

We then employed a within-subject design to assess effects of CNO in cocaine CPP. Following CPP training, a baseline cocaine place preference score was first established on WD1 for each mouse which was then randomly assigned to two groups. One group was injected CNO (1 mg/kg, i.p.) to activate the hM3d(Gq) in IL-NAcSh projection neurons while another group was injected with saline (vehicle control) 30 min prior to CPP test on WD2 and WD15. Administration of saline did not affect CPP on WD2 or WD15 (F_2,26_ = 0.84, *P* = 0.44, one-way repeated measures ANOVA followed by Tukey’s post hoc test; Fig. 5h). In contrast, administration of CNO significantly attenuated cocaine CPP on both WD2 and WD15 (F_2,30_ = 5.09, *P* = 0.012, one-way repeated measures ANOVA followed by Tukey’s post hoc test; Fig. 5h). These results indicate that reduced excitability of IL-NAcSh neurons causally contributed to the cocaine memory acquisition (WD2) and retention (WD15).

## Discussion

In this study, we characterize the intrinsic neurophysiological properties of two adjacent descending mPFC projections to the NAc, the IL-NAcSh and PL-NAcCo circuits, which play prominent yet differential roles in drug seeking, relapse, and extinction. We show that cocaine CPP elicits prolonged hypoexcitability in IL-NAcSh deep layer pyramidal neurons, but not in PL-NAcCo deep layer pyramidal neurons, thus uncovering a PFC subregion-dependent, projection-specific intrinsic plasticity induced by chronic drug experiences. We further demonstrate that chemogenetically restoring the IL-NAcSh neuronal excitability abolishes cocaine-conditioned place memories, suggesting that the IL-NAcSh circuit normally serves to suppress drug associated memories. Our studies provide new insights into how different PFC top-down circuits undergo differential neuroadaptations in response to drug experiences and their roles in the regulation of drug associated memories.

The intrinsic properties of PFC neurons vary depending on their subregions, layer locations and projections. In the rat mPFC, pyramidal neurons in L2/3 were found to be more hyperpolarized and less excitable and possess smaller I_h_ than in L5 in both the PL and IL, and neurons in the IL appear to be more excitable than the PL counterparts due to a lower spike threshold and higher input resistance [39] or a lower rheobase [25]. Within L5 pyramidal neurons of the mPFC, the dopamine D1 receptor-expressing neurons were reported to show lower I_h_ than those expressing D2 receptors that have thick apical tufts, prominent I_h_, and subcortical projections, thus exhibiting different firing patterns [38, 41]. Here, we characterized and directly compared the intrinsic properties of projection-specific mPFC-NAc L5 pyramidal neurons, revealing IL-NAcSh L5 neurons have larger I_h_-mediated rebound potentials, higher excitability, and quantitatively different frequency adaptation in their firing. Whereas the functional significance of these neurophysiological differences remains to be fully investigated, they undergo differential neuroadaptations following cocaine experiences and likely play integral roles in supporting the unique functions of these subregions and circuits in cognitive processes and behaviors.

Due to its importance in drug seeking and extinction behaviors, the mPFC has been investigated in a number of rodent drug-induced intrinsic plasticity studies. Most studies have focused on the PL, the dorsal mPFC, and reported that chronic cocaine exposures (including non-contingent administrations, self-administrations, and place conditioning) increase membrane excitability of deep layer pyramidal neurons in rats with or without withdrawal [23–27]. In contrast, few studies have investigated how drugs of abuse affect the IL, the ventral mPFC. In rats, excitability in IL neurons following extinction from cocaine self-administration was significantly reduced, characterized by fewer depolarization-evoked spikes without overt changes in passive and active membrane properties [25]. An in vivo Ca2+ imaging study in rats self-administering cocaine has also reported decreased activity and spatial selectivity of NAc-projecting IL neurons around the time of cocaine seeking, which were reduced following a drug-free period [15], but the underlying intrinsic and ionic mechanisms were not investigated. Here, we have directly examined the intrinsic active and passive membrane properties of IL-NAcSh L5 pyramidal neurons in mice and demonstrate their long-lasting hypoexcitability following a period of cocaine place conditioning. The intrinsic excitability of PL and IL neurons has also been reported to increase or decrease following exposures to other drugs of abuse, including heroin [42], ethanol [43, 44], and toluene [40].

The cocaine-elicited hypoexcitability in IL-NAcSh L5 neurons is characterized by reduced spiking, higher rheobase, lower input resistance, increased fAHP, modest alteration of AP waveform, and a substantial loss of spike frequency adaptation. Given that these various passive and active membrane properties are differentially mediated by different ion channels, complex ionic mechanisms are likely involved in the loss of excitability in IL-NAcSh neurons. Specifically, neuroadaptations to voltage-gated ion channels (regulating AP waveform and threshold [45]), Ca2+ activated K+ channels (regulating AHPs and spike frequency adaptations [34, 35]), subthreshold Kv7/KCNQ/M channels (regulating AHPs and spike frequency adaptations [27, 46]), and voltage-independent subthreshold leakage/background K+ currents (regulating input resistance [47, 48]) may co-occur following cocaine experiences. Indeed, some of these channels have been shown to undergo neuroadaptations following cocaine exposures in other regions of the reward circuits [21, 27, 49]). HCN channels, which undergo neuroadaptations associated with addiction in other reward regions [50–52], appear not affected here based on unaltered sag and ADP following cocaine CPP. Finally, drug induced intrinsic plasticity may occur in a circuit specific manner, as our results show that PL-NAcCo projection is not similarly affected as is IL-NAcSh projection by cocaine CPP.

The IL, via its descending projections to the amygdala, plays critical roles in the extinction of fear conditioning [53–55]. Analogous to this prefrontal fear extinction circuit, the IL-NAcSh projection regulates extinction of drug seeking behavior and has been proposed as a bona fide anti-relapse circuit [5, 7]. The IL-NAcSh circuit may inhibit drug seeking by suppressing the motivation to seek drugs and/or by extinguishing drug-associated memories, but distinguishing these mechanisms has been less conclusive. Here, using a drug-cue associative memory model, we demonstrate that chemogenetic enhancement of IL-NAcSh neuron excitability, which had been reduced by prior cocaine experiences, attenuates both the acquisition and retention of cocaine conditioned place preference memories. Our study provides strong support for the hypothesis that NAcSh projecting IL output neurons encode information that is necessary to drive the extinction of drug memories.

In conclusion, we show that the IL-NAcSh circuit excitability is susceptible to cocaine exposure and exhibits long-lasting hypoexcitability following cocaine experiences, and this hypoexcitability contributes to the emergence and maintenance of cocaine CPP memory. This cocaine-induced intrinsic plasticity is circuit-dependent as the adjacent PL-NAcCo circuit, whose intrinsic neuronal excitability has been shown to maintain a cocaine CPP memory during retrieval [26], shows a marginal hyperexcitability following a 15 day cocaine withdrawal. Recent studies have also indicated that these PFC-NAc circuits undergo differential synaptic remodeling processes following cocaine self-administration, contributing to incubation of cocaine craving [11]. Drug induced synaptic and intrinsic plasticity may mediate enhanced motivation, established drug memory, and normal inhibitory control of reward behaviors, perpetuating compulsive drug seeking and taking behaviors in addiction.

## ACKNOWLEDGEMENTS

We thank members of the Yao laboratory for critiques and comments. This work was supported by NIH Grants R01DA032283 (to W.-D.Y.).

