## Supplemental figure legends for "A Prefrontal Cortex-Nucleus Accumbens Circuit Attenuates Cocaine-conditioned Place Preference Memories"

**Figure S1**. Spike number-current injection relationships and firing frequency adaptation in individual IL-NAcSh and PL-NAcCo L5 neurons. (**a1, b1**), Number of AP spikes vs. step currents injected in 18 IL-NAcSh (**a1**) and 19 PL-NAcCo (**b1**) L5 pyramidal neurons. (**a2, b2**), firing frequency adaptation in response to 200 pA step current injection in 18 IL-NAcSh (**a2**) and 19 PL-NAcCo (**b2**) L5 pyramidal neurons. Individual cells are color coded.

**Figure S2**. Spike frequency adaptations in individual IL-NAcSh L5 PNs after cocaine withdrawal. (**a**) Firing frequency adaptations in individual neurons from saline mice. (**b**) Firing frequency adaptations in individual neurons form cocaine CPP mice following 1 day withdrawal. (**c**) Firing frequency adaptations in individual neurons form cocaine CPP mice following 15-day withdrawal. APs were evoked by a 200 pA step current injection. Individual cells are color coded.

**Figure S3**. Spike frequency adaptations in individual PL-NAcCo L5 PNs after cocaine withdrawal. Firing frequency adaptations in individual neurons from saline (**a**) and cocaine CPP mice (**b**) after 15-day withdrawal are shown. APs were evoked by a 200 pA step current injection. Individual cells are color coded.
