## Supplementary figures and images for "A Prefrontal Cortex-Nucleus Accumbens Circuit Attenuates Cocaine-conditioned Place Preference Memories"

### Supplemental figure 1

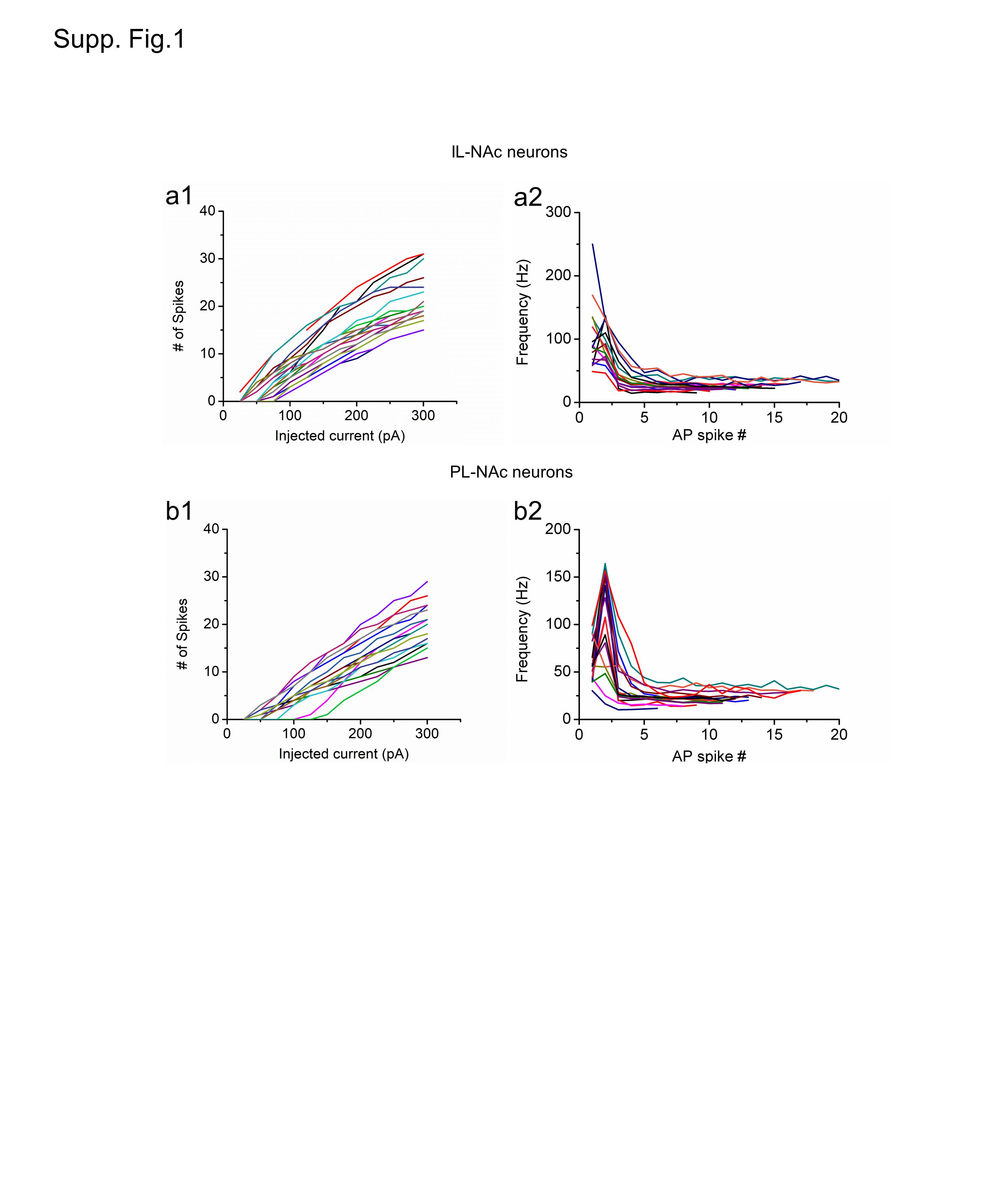

### Supplemental figure 2

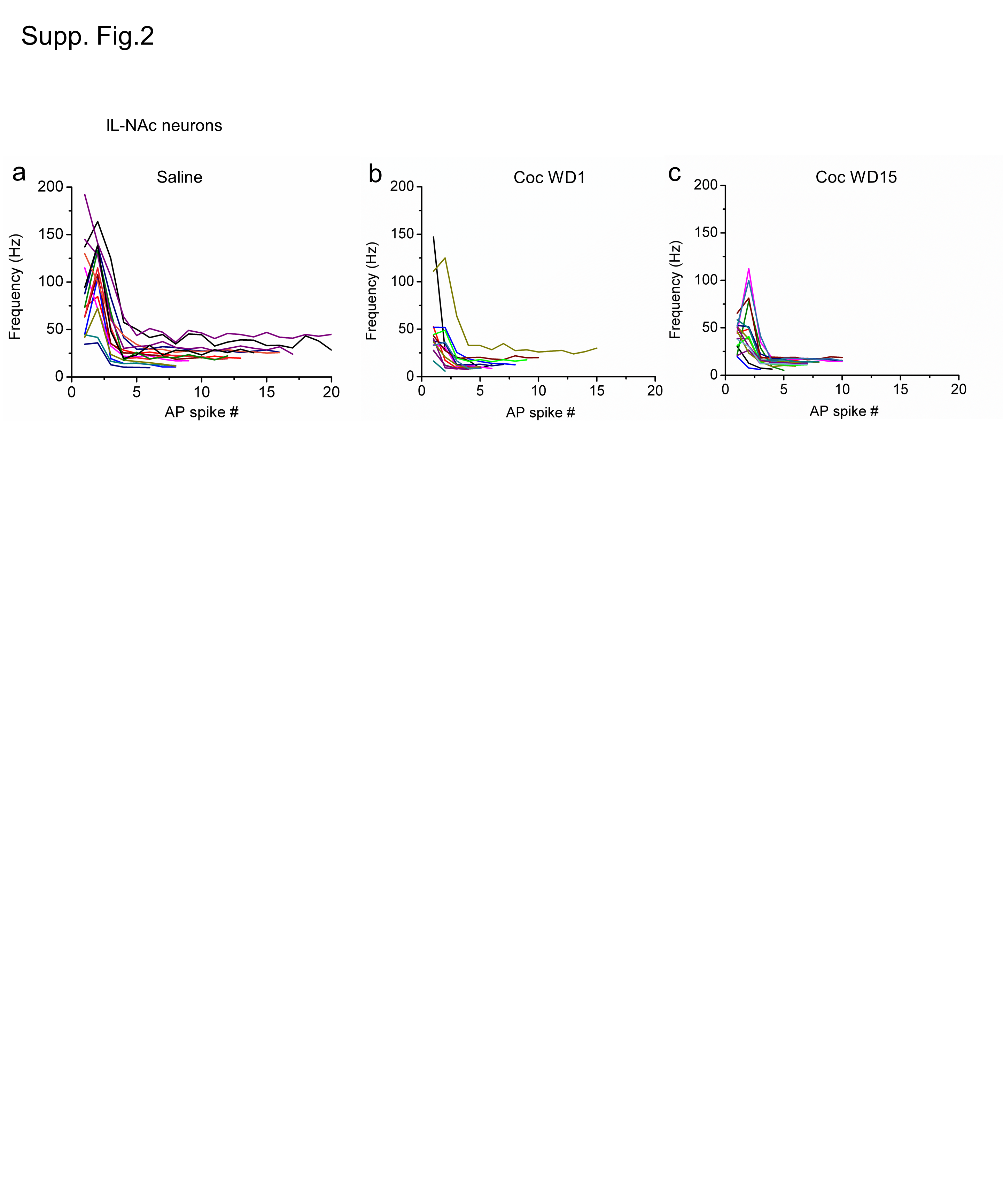

### Supplemental figure 3

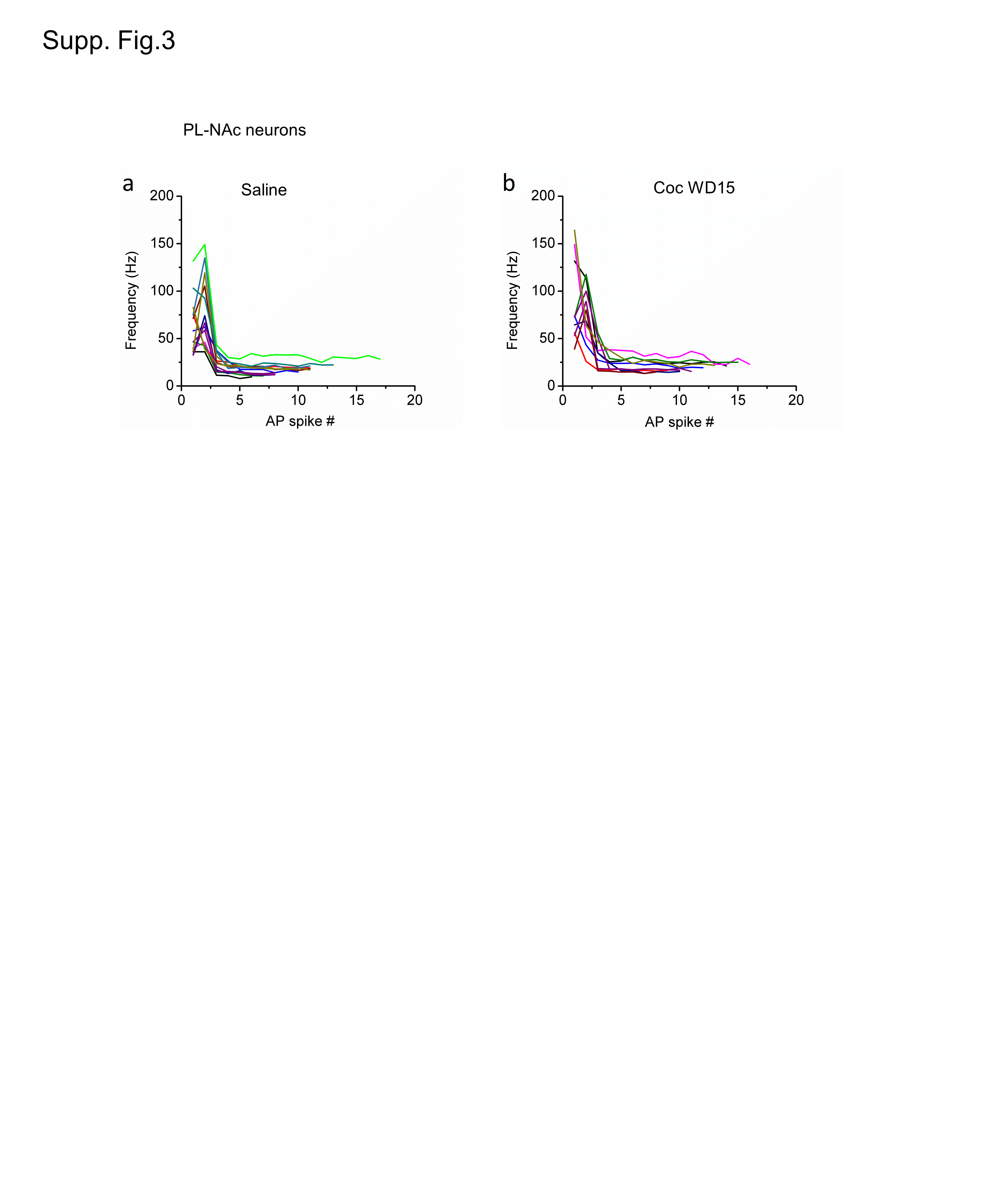
